# Structure, immunogenicity, and conformation-dependent receptor binding of the post-fusion human metapneumovirus F protein

**DOI:** 10.1101/2021.05.19.444868

**Authors:** Jiachen Huang, Pradeep Chopra, Lin Liu, Tamas Nagy, Jackelyn Murray, Ralph A. Tripp, Geert-Jan Boons, Jarrod J. Mousa

## Abstract

Human metapneumovirus (hMPV) is an important cause of acute viral respiratory infection. As the only target of neutralizing antibodies, the hMPV fusion (F) protein has been a major focus for vaccine development and targeting by drugs and monoclonal antibodies (mAbs). While X-ray structures of trimeric pre-fusion and post-fusion hMPV F proteins from genotype A, and monomeric pre-fusion hMPV F protein from genotype B have been determined, structural data for the post-fusion conformation for genotype B is lacking. We determined the crystal structure of this protein and compared the structural differences of post-fusion hMPV F between hMPV A and B genotypes. We also assessed the receptor binding properties of the hMPV F protein to heparan sulfate. A library of heparan sulfate (HS) oligomers was used to verify the HS binding activity of hMPV F, and several compounds showed binding to predominantly pre-fusion hMPV F, but had limited binding to post-fusion hMPV F. Furthermore, mAbs to antigenic sites III and the 66-87 intratrimeric epitope block heparan binding. In addition, we evaluated the efficacy of post-fusion hMPV B2 F protein as a vaccine candidate in BALB/c mice. Mice immunized with hMPV B2 post-fusion F protein showed a balanced Th1/Th2 immune response and generated neutralizing antibodies against both subgroup A2 and B2 hMPV strains, which protected the mice from hMPV challenge. Antibody competition analysis revealed the antibodies generated by immunization target two known antigenic sites (III and IV) on hMPV F. Overall, this study provides new characteristics of the hMPV F protein informative for vaccine and therapy development.

**Importance:** Human metapneumovirus (hMPV) is an important cause of viral respiratory disease. In this paper, we report the X-ray crystal structure of the hMPV fusion (F) protein in the post-fusion conformation from genotype B. We also assessed binding of the hMPV F protein to heparin and heparan sulfate, a previously reported receptor for the hMPV F protein. Furthermore, we determined the immunogenicity and protective efficacy of post-fusion hMPV B2 F protein, which is the first study using a homogenous conformation of the protein. Antibodies generated in response to vaccination give a balanced TH1/TH2 response and target two previously discovered neutralizing epitopes.

## Introduction

Human metapneumovirus (hMPV) is a negative-sense single-stranded enveloped RNA virus in the family of Pneumoviridae. There are two circulating genotypes of hMPV (A and B), which are further divided into four subgroups, A1, A2, B1, and B2, based on the sequence variability of the surface proteins (1). hMPV is one of the major causes of respiratory infections affecting infants and children under 5 years of age, and accounts for 6-40% cases of acute respiratory infections in hospitalized and outpatient children (2). Serological studies showed that almost all people are exposed to hMPV by age 5 (3–5), and reinfections can happen throughout life. Premature infants, the elderly and immunocompromised patients are at high risk of severe diseases caused by hMPV infection (2, 6). However, there are no licensed vaccines or specific treatments available for hMPV infection.

The fusion (F) protein of hMPV is highly conserved among hMPV subgroups, and it shares similar structural topology and approximately 30% amino acid sequence homology with the respiratory syncytial virus (RSV) F protein. In addition, the hMPV F protein, as with all Pneumovirus and Paramyxovirus F proteins, plays an indispensable role in viral infection. hMPV F belongs to the family of class I viral fusion proteins that mediate the fusion of viral envelope and cell membrane during infection. hMPV F is first synthesized as a polypeptide precursor, F0, and is then cleaved by an unknown enzyme to generate a F1-F2 heterodimer connected by disulfide bonds, which form the mature trimeric pre-fusion structure. The pre-fusion conformation of hMPV F is meta-stable and undergoes conformational rearrangement to the post-fusion state during the process of membrane fusion (7). In addition, hMPV F is involved in virus attachment and receptor binding. Heparan sulfate (HS), a glycosaminoglycan that is ubiquitously expressed on the membrane surface of all animal tissues, has been hypothesized to be a receptor for the hMPV F protein (8). HS blocks has been shown to block hMPV from infecting human lung cells and airway tissues *in vitro* (9). Integrin α_5_β1 is another potential cellular receptor for hMPV F, as the hMPV F protein has an Arg-Gly-Asp (RGD) binding motif (10, 11). Function-blocking monoclonal antibodies (mAbs) targeting α_5_β1 integrin, siRNA targeting α_5_ or β1, and EDTA all disrupt hMPV infection (12). Mutagenesis of the RGD motif inhibits cell-cell fusion, and mutant viruses have impaired growth *in vitro* and *in vivo* (13). However, there is still no evidence to show direct interactions between the hMPV F protein and these potential host receptors, and it is unclear whether hMPV F-specific mAbs can block receptor binding of hMPV F.

As the only target of neutralizing antibodies (14), hMPV F has been stabilized in both pre-fusion and post-fusion conformations to facilitate recombinant expression and vaccine development (15, 16). The majority of hMPV F-specific human antibodies bind hMPV F in both pre-fusion and post-fusion conformations (15, 17), while pre-fusion RSV F is preferred by neutralizing human antibodies (18). Like formalin-inactivated (FI)-RSV vaccines that induced aberrant immune responses and lead to enhanced respiratory disease in children after natural RSV reinfection (19– 21), FI-hMPV and heat-inactivated hMPV vaccines also caused enhanced disease following viral infection in mice, cotton rats, and macaques potentially due to an abnormal Th2 immune response that leads to increased cytokine levels and lung inflammation (22, 23). Other forms of hMPV vaccines have also been explored in recent years. A recombinant live attenuated hMPV vaccine was tested in a phase I clinical trial in adults and children, but the vaccine was over attenuated and failed to efficiently infect hMPV-seronegative children (24). Several viral vector-based or virus-like particle (VLP)-based hMPV vaccine candidates have also been evaluated in animal models and pre-clinical studies (14, 25, 26), which showed encouraging results. A bivalent fusion protein-based hMPV/PIV3 mRNA vaccine is currently under Phase I clinical trials. With proper adjuvant, hMPV F-based subunit vaccines can induce protective immunity without enhancement of disease in cotton rats and non-human primates (27, 28), indicating hMPV F is a promising vaccine candidate. The crystal structure of post-fusion hMPV A1 F has been solved and it can induce neutralizing antibodies after one immunization with CpG adjuvant in mice (16). However, there remain several uncertainties regarding hMPV F vaccination, including the potential of post-fusion hMPV F immunization to prevent viral replication, and the antigenic epitopes on the hMPV F protein targeted by B cells.

To further explore the structural, immunological, and receptor binding properties of hMPV F, we determined the crystal structure of post-fusion hMPV F protein from a genotype B strain. In addition, we determined the capacity of hMPV F to bind HS, and determined that neutralizing hMPV F mAbs can block HS binding. Furthermore, the immunological properties of the protein were assessed by vaccination and challenge studies.

## Methods

### Ethics statement

All animal studies were approved by the University of Georgia Institutional Animal Care and Use Committee.

### Production and purification of recombinant hMPV F proteins

Plasmids encoding cDNA for hMPV 130-BV F and hMPV B2 F proteins were synthesized (GenScript) and cloned into the pcDNA3.1+ vector as previously described (29, 30). The plasmids were expanded by transformation in Escherichia coli DH5α cells with 100 µg/mL of ampicillin (Thermo Fisher Scientific) for selection. Plasmids were purified using the E.Z.N.A. plasmid maxi kit (Omega BioTek), according to the manufacturer’s protocol. The stable cell line that expresses the hMPV B2 F protein was generated as previously described (30). For protein expression and purification, the stable cell lines were expanded in 500 mL of Freestyle293 medium supplemented with G418 at 1 × 106 cells/mL. After 5 to 7 days, recombinant protein was purified from the filtered culture supernatant using HisTrap Excel columns (GE Healthcare Life Sciences). Each column was stored in 20% ethanol and washed with 5 column volumes (CV) of wash buffer (20 mM Tris pH 7.5, 500 mM NaCl, and 20 mM imidazole) before loading samples onto the column. After sample application, columns were washed with 10 CV of wash buffer. Proteins were eluted from the column with 6 CV of elution buffer (20 mM Tris pH 7.5, 500 mM NaCl, and 250 mM imidazole). Proteins were concentrated and buffer exchanged into phosphate buffered saline (PBS) using Amicon Ultra-15 centrifugal filter units with a 30 kDa cutoff (Millipore Sigma).

### Trypsinization of hMPV F

In order to generate homogeneous cleaved trimeric hMPV F protein, trypsin-tosylsulfonyl phenylalanyl chloromethyl ketone (TPCK) (Thermo Scientific) was dissolved in double-distilled water (ddH2O) at 2 mg/mL. Purified hMPV B2 F was incubated with 5 TAME (p-toluene-sulfonyl-L-arginine methyl ester) units/mg of trypsin-TPCK (L-1-tosylamido-2-phenylethyl chloromethyl ketone) for 1 h at 37 °C. Trimeric hMPV B2 F protein was purified from the digestion reaction mixture by size exclusion chromatography on a Superdex S200, 16/600 column (GE Healthcare Life Sciences) in column buffer (50 mM Tris pH 7.5, and 100 mM NaCl). The trimeric hMPV F protein was identified by a shift in the elution profile from monomeric hMPV B2 F protein. The fractions containing the trimers and monomers were concentrated using 30 kDa Spin-X UF concentrators (Corning). To obtain homogenous post-fusion hMPV F, trimeric trypsinized hMPV F was heated at 55 °C for 20 minutes to induce conversion of remaining pre-fusion hMPV F proteins to the post-fusion conformation.

### Negative-stain electron microscopy

All samples were purified by size exclusion chromatography on a Superdex S200, 16/600 column (GE Healthcare Life Sciences) in column buffer before they were applied on grids. Carbon-coated copper grids (Electron Microscopy Sciences) were overlaid with 5 μl of protein solutions (10 μg/mL) for 3 min. The grid was washed in water twice and then stained with 0.75% uranyl formate for 1 min. Negative-stain electron micrographs were acquired using a JEOL JEM1011 transmission electron microscope equipped with a high-contrast 2K-by-2K AMT midmount digital camera.

### Human mAb production and purification

For recombinant mAbs, plasmids encoding cDNAs for heavy and light chain sequences of 101F (31), MPE8 (31), and DS7 (32) were synthesized (GenScript), and were cloned into vectors encoding human IgG1 and lambda or kappa light chain constant regions, respectively. mAbs were obtained by transfection of plasmids into Freestyle HEK293F cells as described previously (30). For hybridoma-derived mAbs, hybridoma cell lines were expanded in serum-free medium (Hybridoma-SFM; Thermo Fisher Scientific). Recombinant cultures from transfection were stopped after 5 to 7 days, hybridoma cultures were stopped after 30 days. Culture supernatants from both approaches were filtered and mAbs were purified from culture supernatants using HiTrap protein G columns (GE Healthcare Life Sciences) according to the manufacturer’s protocol.

### Crystallization and structure determination of the trimeric post-fusion hMPV B2 F

Purified trypsinized post-fusion hMPV B2 F protein was subjected to size exclusion chromatography (S200, 16/300, GE Healthcare Life Sciences) in 50 mM Tris pH 7.5, 100 mM NaCl. The fractions containing the trimeric hMPV F protein were concentrated to 12 mg/mL and crystallization trials were prepared on a TTP LabTech Mosquito Robot in sitting-drop MRC-2 plates (Hampton Research) using several commercially available crystallization screens. Crystals were obtained in the Index HT (Hampton Research) in condition A6 (0.1 M Tris pH 8.5, 2.0 M Ammonium sulfate). Crystals were harvested and cryo-protected with 30% glycerol in the mother liquor before being flash frozen in liquid nitrogen. X-ray diffraction data were collected at the Advanced Photon Source SER-CAT beamLine 21-ID-D. Data were indexed and scaled using XDS (33). A molecular replacement solution was obtained in Phaser (34) using the post-fusion hMPV A1 F structure (PDB 5L1X). The crystal structure was completed by manually building in COOT (35) followed by subsequent rounds of manual rebuilding and refinement in Phenix. The data collection and refinement statistics are shown in **Table 1**.

**Table 1.** Summary of X-ray diffraction and model statistics.

| **Table 1. Summary of X-ray diffraction and model statistics.** |  |
| --- | --- |
| Wavelength | 1 Å |
| Resolution range | 49.65 - 3.1 (3.21 - 3.1) |
| Space group | P 21 2 21 |
| Unit cell | 114.6 128.1 431.7<br>90 90 90 |
| Total reflections | 232377 (22844) |
| Unique reflections | 116215 (11420) |
| Multiplicity | 2.0 (2.0) |
| Completeness (%) | 99.9 (99.9) |
| Mean I/sigma(I) | 7.5 (1.8) |
| Wilson B-factor | 47.4 |
| R-merge | 0.114 (0.482) |
| R-meas | 0.161 (0.681) |
| R-pim | 0.114 (0.482) |
| CC1/2 | 0.974 (0.650) |
| CC* | 0.993 (0.888) |
| Reflections used in refinement | 116149 (11420) |
| Reflections used for R-free | 11536 (1131) |
| R-work | 0.230 (0.322) |
| R-free | 0.277 (0.368) |
| CC (work) | 0.929 (0.764) |
| CC (free) | 0.893 (0.674) |
| Number of non-hydrogen atoms | 19737 |
| macromolecules | 19362 |
| ligands | 252 |
| solvent | 123 |
| Protein residues | 2538 |
| RMS(bonds) | 0.012 |
| RMS(angles) | 1.34 |
| Ramachandran favored (%) | 90.53 |
| Ramachandran allowed (%) | 9.23 |
| Ramachandran outliers (%) | 0.24 |
| Rotamer outliers (%) | 0.32 |
| Clashscore | 21.4 |
| Average B-factor | 58.8 |
| macromolecules | 58.6 |
| ligands | 89.2 |
| solvent | 30.0 |

### HS microarray printing and screening

All compounds were printed on NHS-ester activated glass slides (NEXTERION® Slide H, Schott Inc.) using a Scienion sciFLEXARRAYER S3 non-contact microarray equipped with a Scienion PDC80 nozzle (Scienion Inc.). Individual samples were dissolved in sodium phosphate buffer (50 µL, 0.225 M, pH 8.5) at a concentration of 100 µM, and were printed in replicates of 10 with spot volume ∼ 400 pL, at 20 °C and 50% humidity. Each slide has 24 subarrays in a 3×8 layout. After printing, slides were incubated in a humidity chamber for 8 hrs and then blocked for 30 min with a 5 mM ethanolamine in a Tris buffer (pH 9.0, 50 mM) at 40 °C. Blocked slides were rinsed with DI water, spun dry, and kept in a desiccator at room temperature for future use. Screening was performed by incubating the slides with a protein solution for 1 hour followed by washing and drying. The buffers used in screening are TSM buffer (TSM, 20 mM Tris·Cl, pH 7.4, 150 mM NaCl, 2 mM CaCl2, and 2 mM MgCl2), TSM binding buffer (TSMBB, TSM buffer with 0.05% Tween-20 and 1% BSA) and TSM washing buffer (TSMWB, TSM buffer with 0.05% Tween-20). A typical washing procedure includes sequentially dipping the glass slide in TSM wash buffer (2 min, containing 0.05 % Tween 20), TSM buffer (2 min) and, water (2 × 2 min), followed by spun dry.

The slides were incubated with His-tagged proteins diluted in TSMBB at different concentrations for 1 hr, followed by washing and incubated with a solution of AlexaFluor® 647 conjugated anti-His antibody (BioLegend #652513, 10 µg/mL). After washing and drying, the slides were scanned using a GenePix 4000B microarray scanner (Molecular Devices) at 635 nm with a resolution of 5 µM. Various gains and PMT values were employed for the scanning to ensure that all the signals were within the linear range of the scanner’s detector and there was no saturation of signals. The images were analyzed using GenePix Pro 7 software (version 7.2.29.2, Molecular Devices). The data were analyzed with a home written Excel macro. The highest and the lowest value of the total fluorescence intensity of the replicates spots were removed, and the remaining values were used to provide the mean value and standard deviation (36, 37).

### Heparin binding assay (SPR)

For the preparation of a heparin sensor chip, CM5 chip was first coated with streptavidin by standard amine coupling using an amine coupling kit (Biacore, GE Healthcare), followed by immobilization of biotin-heparin (38). Briefly, the surface was activated using freshly mixed N-hydroxysuccinimide (NHS; 100 mM) and 1-(3-dimethylaminopropyl)-ethylcarbodiimide (EDC; 391 mM) (1/1, v/v) in water. Next, streptavidin (50 μg/mL, Invitrogen) in aqueous NaOAc (10 mM, pH 4.5) was passed over the chip surface until a ligand density of approximately 2000 RU was achieved. The remaining NHS-activated esters were quenched by aqueous ethanolamine (1.0 M, pH 8.5). Next, biotin-heparin (50 μg/mL) was passed over one of the flow channels at a flow rate of 10 μL/min for 30 s resulting in a response of 83 RU. Next, the reference and modified flow cells were washed with three consecutive injections of 60 s with 1.0 M NaCl. Phosphate-buffered saline (PBS, pH 7.4) was used as the running buffer for the immobilization, kinetic studies and inhibition studies. Analytes were dissolved in running buffer and a flow rate of 30 μL/min was employed for association (180 s) and dissociation (600 s) at a constant temperature of 25 °C. A 30 s injection of 1.0 M NaCl at a flow rate of 30 μL/min was used for regeneration and to achieve prior baseline status. Using Biacore T100 evaluation software, the response curves of various analyte concentrations were globally fitted to the two-state binding model.

### Growth of hMPV

hMPV B2 strain TN/93-32 and hMPV A2 strain CAN/97-83 viruses were grown in LLC-MK2 cells (ATCC) as previously described (30). Briefly, cells were grown to 80% confluence in 225 cm^2^ flasks in Opti-MEM supplemented with 2% FBS. For virus infection, cells were washed twice with Dulbecco’s phosphate-buffered saline (DPBS; Corning) and then infected with hMPV at 1:100 MOI, supplemented with 5 µg/mL trypsin-EDTA and 100 µg/mL CaCl2. Cells were incubated for five days, the medium was removed from the flask, and 5 mL of cold 25% (wt/vol) sterile-filtered sucrose was added to the flask. The flask was transferred to −80 °C until the solution was frozen, then moved to thaw at 4 °C, followed by another freeze-thaw cycle. Cell lysates were scraped and transferred to a sterile tube and centrifuged at 1,100 rpm for 5 min to remove cell debris. The clarified aliquoted of supernatant containing hMPV was flash frozen and titered for later use.

### Mice immunization and hMPV challenge

BALB/c mice (6-8 weeks old, Charles River Laboratories) were immunized in a prime-boost-boost regimen with post-fusion hMPV B2 F protein (50 µg protein/mouse) in a water in oil emulsion with Titermax Gold adjuvant (Sigma-Aldrich) via subcutaneous (SC) route into the loose skin over the neck, while mice in control groups were immunized with PBS + TiterMax Gold adjuvant emulsion. Two weeks after prime, the mice were boosted with the same amount of the emulsion, and a second boost without adjuvant was performed two weeks after the first boost. For the challenge study, BALB/c mice (6-8 weeks old, The Jackson Laboratory) were vaccinated with the same dosage as above, but only boosted once with adjuvant 4 weeks after the prime. Two weeks after the boost, mice were intranasally challenged with hMPV TN/93-32 (∼5×10^5^ pfu per mice). Mice were sacrificed 5 days post-challenge, and lungs were collected for virus titration and histological analysis.

### Immunostaining of hMPV plaques

For serum neutralization assay, heat inactivated mouse serum was serial diluted and incubated 1:1 with a suspension of hMPV for 1 hr at room temperature. Following this, LLC-MK2 cells in 24 well plates were inoculated with the serum-virus mixture (50 µl per well) for 1 hr and rocked at room temperature. Cells were then overlaid with 1 mL of 0.75% methylcellulose dissolved in Opti-MEM supplemented with 5 µg/mL trypsin-EDTA and 100 µg/mL CaCl2. Cells were incubated for 4 days, after which the cells were fixed with 10% neutral buffered formalin. The cell monolayers were then blocked with blocking buffer (2% Blotting-Grade Blocker [Bio-Rad] supplemented with 2% goat serum [Gibco] in PBS-Tween) for 1 hr. The plates were washed with water, and 200 µl of MPV364 (29) primary antibody (1 µg/mL diluted in blocking buffer) was added to each well, and the plates were incubated for 1 hr. The plates were then washed three times with water, after which 200 µl of goat anti-human IgG-horseradish peroxidase (HRP) secondary antibody (catalog number 5220-0286; SeraCare) diluted 1:2,000 in blocking buffer was added to each well for 1 hr. Plates were then washed five times with water, and 200 µl of TrueBlue peroxidase substrate (SeraCare) was added to each well. Plates were incubated until plaques were clearly visible. Plaques were counted by hand under a stereomicroscope and compared to a virus-only control, and the data were analyzed in GraphPad Prism using a nonlinear regression curve fit and the log(inhibitor)-versus-response function. To determine the viral load in lungs of challenged mice, the lungs were collected after euthanasia and homogenized with gentleMACS M Tubes (Miltenyi Biotec) in 1 mL Opti-MEM, followed by centrifugation at 300g for 10 minutes to pellet the tissue debris. The supernatant was aliquoted and serial diluted with Opti-MEM, then plated on LLC-MK2 cells in 24 well plates (100 uL per well). After a 1 hr incubation at 37 °C, cells were then overlaid with 1 mL of 0.75% methylcellulose dissolved in Opti-MEM supplemented with 5 µg/mL trypsin-EDTA, 100 µg/mL CaCl2, and 1X Antibiotic-Antimycotic (Gibco). After 5-7 days, the plaques were stained in the same way described above.

### Enzyme-linked immunosorbent assay for binding to hMPV F proteins

For recombinant protein capture ELISAs, 384-well plates (Greiner Bio-One) were coated with 2 μg/mL of antigen in PBS overnight at 4 °C. Following this, plates were washed once with water before blocking for 1 hour with blocking buffer. Primary mAbs or serial dilutions of mouse serum were applied to wells for 1 hour following three washes with water. Plates were washed with water three times before applying 25 μL of secondary antibody (goat anti-mouse IgG Fc, BioLegend) at a dilution of 1:4,000 in blocking solution. After incubation for 1 hour, the plates were washed five times with PBS-Tween, and 25 μL of a PNPP (p-nitrophenyl phosphate) substrate solution (1 mg/mL PNPP in 1 M Tris base) was added to each well. The plates were incubated at room temperature for 1 hr before reading the optical density at 405 nm on a BioTek plate reader. Data was analyzed in GraphPad Prism using a nonlinear regression curve fit and the log(agonist)-versus-response function to calculate the binding EC50 values. Antibody titers were calculated from the highest dilution of a serum sample that produced OD readings of >0.3 above the background readings and were shown in a log10 scale as previously described (39). For competition ELISAs, 384 well plates were coated with 2 μg/mL of trypsinized hMPV B2 F monomer and trimer in PBS overnight at 4 °C and then blocked for 1 hr. Serial dilutions of mouse serum were premixed with equal volume of competing human mAbs (1 μg/mL) in blocking buffer and then applied to wells as primary antibodies for 1 hr. Mouse IgG and human IgG were detected with separate secondary antibodies (goat anti-human IgG Fc, goat anti-mouse IgG Fc, BioLegend) at a dilution of 1:4,000 in blocking buffer and PNPP as described above.

### Pulmonary histopathological analysis

After euthanasia, the lungs were collected, expanded through the trachea with 10% neutral-buffered formalin (NBF) and immersion-fixed with 10% NBF. Fixed lungs were embedded in paraffin, sectioned at 4.0 µm thickness, mounted on positively charged glass slides, stained with hematoxylin-eosin (HE), and coverslipped. Histological sections were evaluated by a board-certified veterinary pathologist. Histopathological scoring was performed according to previously established histopathologic criteria (23). Briefly, peribronchiolitis, perivasculitis, interstitial pneumonitis, and alveolitis were reviewed and scored on a scale of 0-4.

## Results

### X-ray crystal structure of post-fusion hMPV B2 F protein

Stabilized post-fusion hMPV A1 F protein has previously been generated by expression in CV-1 cells by removal of the hMPV F fusion peptide (residues 103–111) to prevent aggregation, replacing the transmembrane domain with the fibritin trimerization domain (Foldon) from T4 bacteriophage, and altering the cleavage site with the second furin-cleavage site of hRSV F (16). We previously reported a post-fusion hMPV B2 F protein by incorporating similar genetic modifications, but this protein was expressed in HEK293F cells (29, 30). Here, we utilized this protein construct to determine the X-ray crystal structure of post-fusion hMPV F from subgroup B2. Recombinantly expressed hMPV B2 F protein was subjected to trypsin digestion to induce cleavage, and both monomeric and trimeric hMPV F were isolated by size exclusion chromatography **(Figure 1A, 1B)**. Limited cleavage was observed in a reducing SDS-PAGE without trypsin cleavage, however, the addition of trypsin results in multiple bands for the hMPV F monomer and a shift from the aggregated post-fusion hMPV F to monomeric hMPV F for the trimeric fraction **(Figure 1A)**. We previously reported that the monomeric hMPV B2 F fraction is in the pre-fusion conformation based on the X-ray crystal structure of hMPV B2 F with the neutralizing mAb MPV458 (30). This trimeric fraction contains a mixture of elongated post-fusion-like and spherical pre-fusion-like particles **(Figure 1C)**. To obtain homogeneous trimeric post-fusion hMPV F, the protein was heated at 55 °C to convert all particles to the post-fusion conformation **(Figure 1D)**. This protein was crystallized and the X-ray crystal structure of this protein was determined to 3.1 Å **(Figure 2, Table 1)**. Three N-linked glycans were observed at residues 57, 172 and 353 on each protomer and residues that are different from A1 F are labeled in red **(Figure 2A)**. The overall structure of post-fusion hMPV B2 F protein highly resembles the reported structure of post-fusion hMPV F A1 F with an RMSD of 0.697 Å **(Figure 2B)**. Approximately 15 amino acid residues on the N-terminus of the F1 domain are missing in the structure, possibly due to the cleavage by trypsin at site Arg129.

**Figure 1.**
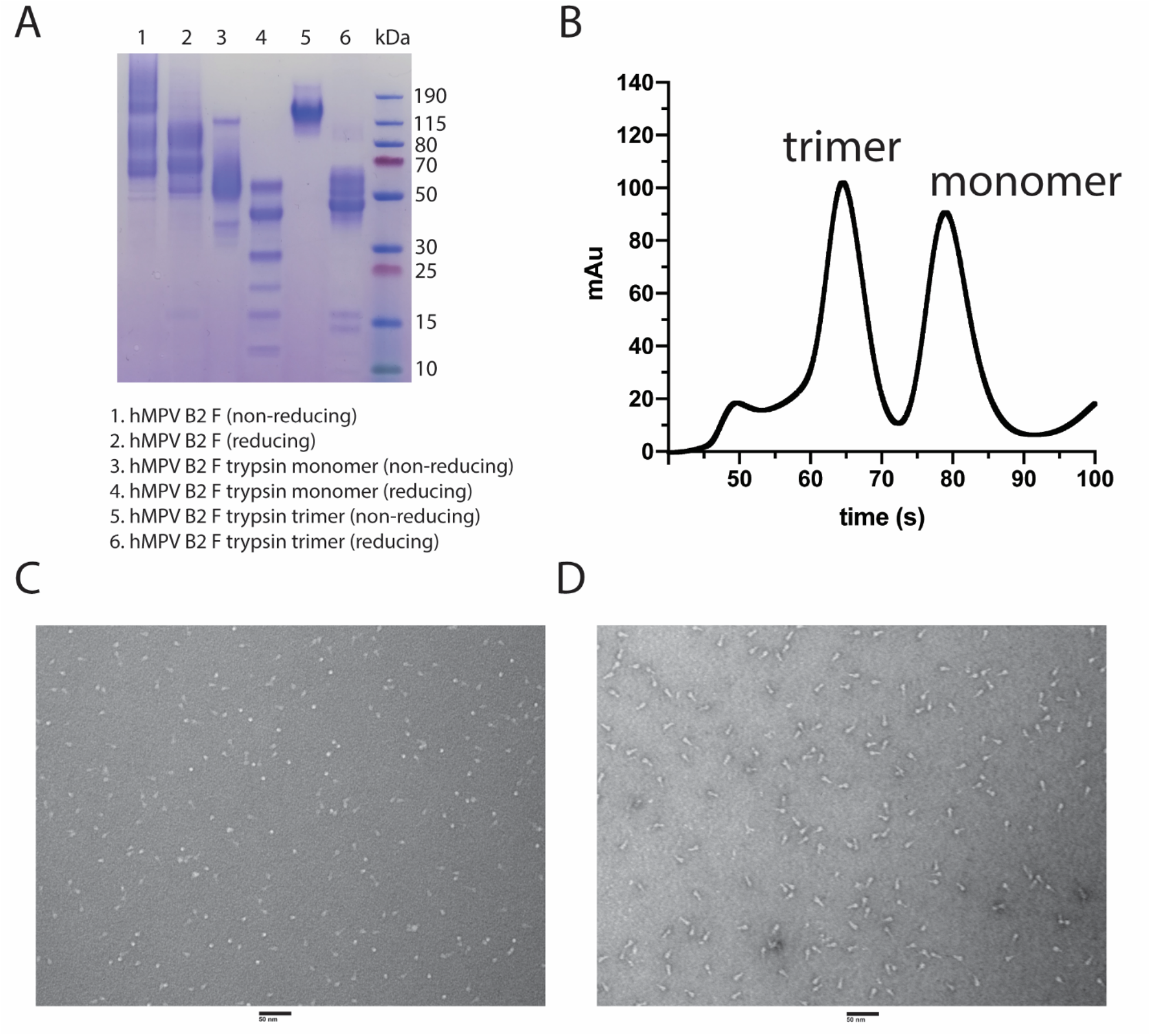
Analysis of the hMPV B2 F protein. (A) SDS-PAGE of hMPV B2 F protein before and after trypsinization. Both monomeric and trimeric fractions were also isolated and analyzed (B) Size exclusion chromatography curve of trypsinized hMPV B2 F protein. Negative-stain electron micrograph of trypsinized hMPV B2 F trimeric protein before (C) and after (D) heating to 55 °C. The scale bar indicates 50 nm.

**Figure 2.**
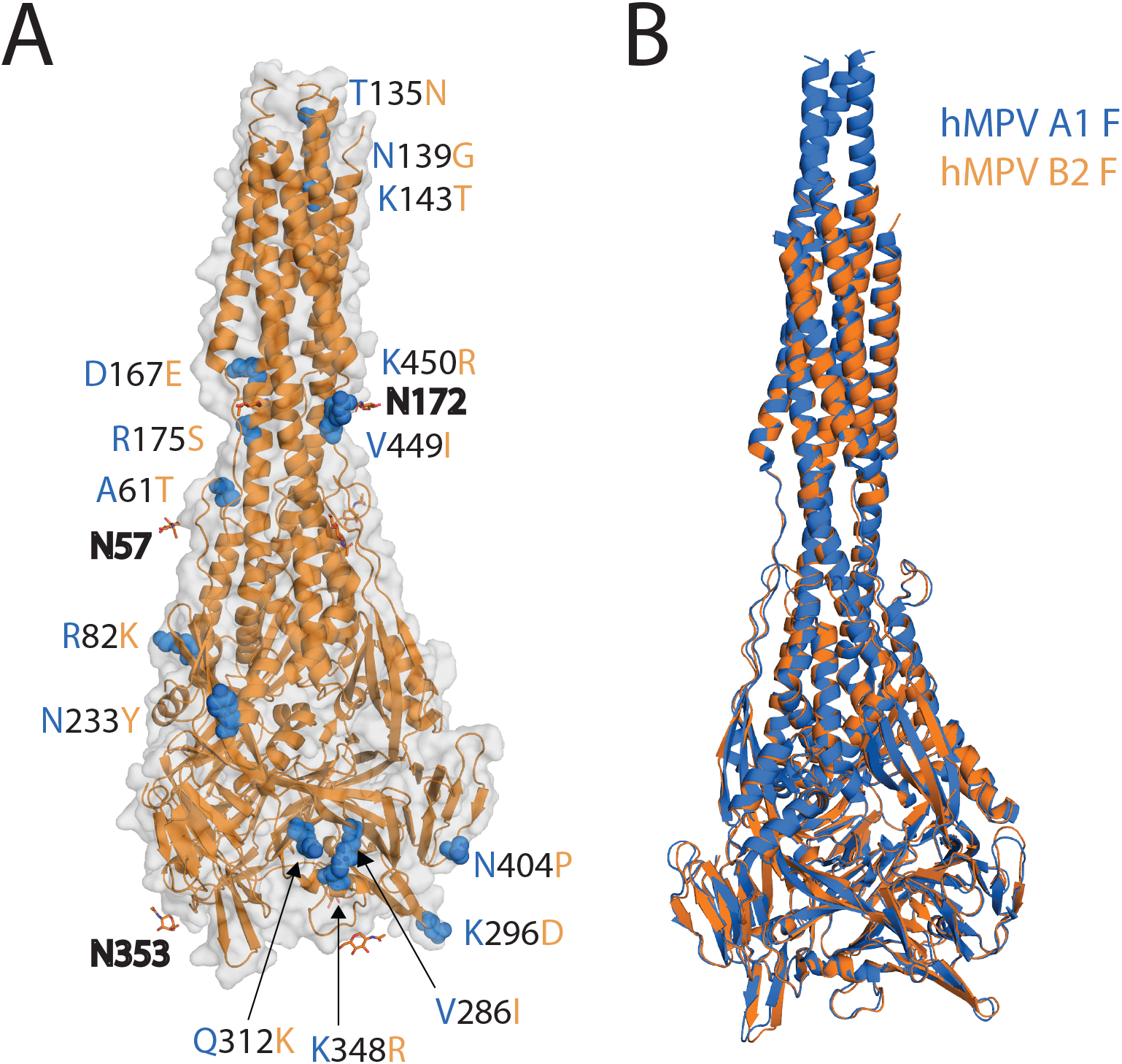
X-ray crystal structure of the post-fusion hMPV B2 F protein. (A) The post-fusion hMPV B2 F protein is displayed. Different residues between hMPV B2 F and A1 F are labeled in red. (B) Overlay of post-fusion hMPV A1 F (cyan, PDB 5L1X) and B2 F (green, PDB 7M0I).

### Conformation-dependence and epitope specificity of heparan sulfate binding by hMPV F

We next sought to probe the receptor binding properties of this protein. Both heparan sulfate and integrin α_5_β1 have been identified as receptors for the hMPV F protein (8, 11). As it is possible that heparan sulfate binding epitopes are not present in the post-fusion conformation, we recombinantly expressed a previously described hMPV F protein with an A185P stabilizing mutation, hMPV 130-BV (15). This protein contains the native hMPV F cleavage site and would not be proteolytically cleaved in HEK293F cells. To determine the conformation of the hMPV 130-BV F protein, we measured the binding of two pre-fusion-specific human mAbs, MPE8 (40), and the recently described MPV465 (30) by ELISA. The epitope for MPE8 on the RSV F protein spans two protomers at antigenic site III (41), and such an epitope would also be expected for hMPV F as this mAb neutralizes both viruses. In contrast, MPV465 binds monomeric hMPV F in the pre-fusion conformation (30). MPE8 had weak binding to monomeric and post-fusion hMPV B2 F, while having 30-fold higher binding to the hMPV 130-BV F protein compared to post-fusion hMPV B2 F **(Figure 3A)**. MPV465 had weak binding to post-fusion hMPV B2 F, yet showed approximately 25-fold and 70-fold higher binding to hMPV 130-BV F and hMPV B2 F monomer, respectively. hMPV 130-BV F has a mixture of pre-fusion and post-fusion-like particles in negative strain electron micrographs **(Figure 3B)**. These data suggest the hMPV 130-BV F protein at least partially mimics the pre-fusion conformation of hMPV F. Binding of heparin to hMPV B2 F protein was examined by surface plasmon resonance, and limited binding over the PBS control was observed for both monomeric (pre-fusion) and trimeric (post-fusion) proteins **(Figure 3C)**. In contrast, heparin binding to hMPV F 130-BV showed an approximately 60-fold improvement, suggesting a trimeric pre-fusion conformation is required for optimal heparin recognition **(Figure 3C)**. A hexa-his peptide **(Figure 3D)** was also tested as a control, to ensure that the his-tag has no contribution to heparin or HS saccharide binding by the protein. Heparin bound to hMPV F 130-BV with a KD of 2.1 nM **(Figure 3E)**, and the binding can be inhibited by unfractionated heparin (UFH) in a dose-dependent manner **(Figure 3F)**. To determine if heparin binding is localized to a single epitope or binds to multiple sites, we assessed binding of heparin to hMPV F 130-BV in the presence of multiple concentrations of mAbs MPE8 and MPV458 (30). In both cases, the binding signal decreased as the concentration of mAb increased **(Figure 3G, 3H)**. We also tested competition with mAbs 101F, and DS7, however, these two mAbs bound nonspecifically to heparin (data not shown). Overall, these data suggest heparin binding to hMPV F occurs optimally to the trimeric pre-fusion conformation, and that binding occurs at or near the MPE8 and MPV458 epitopes.

**Figure 3.**
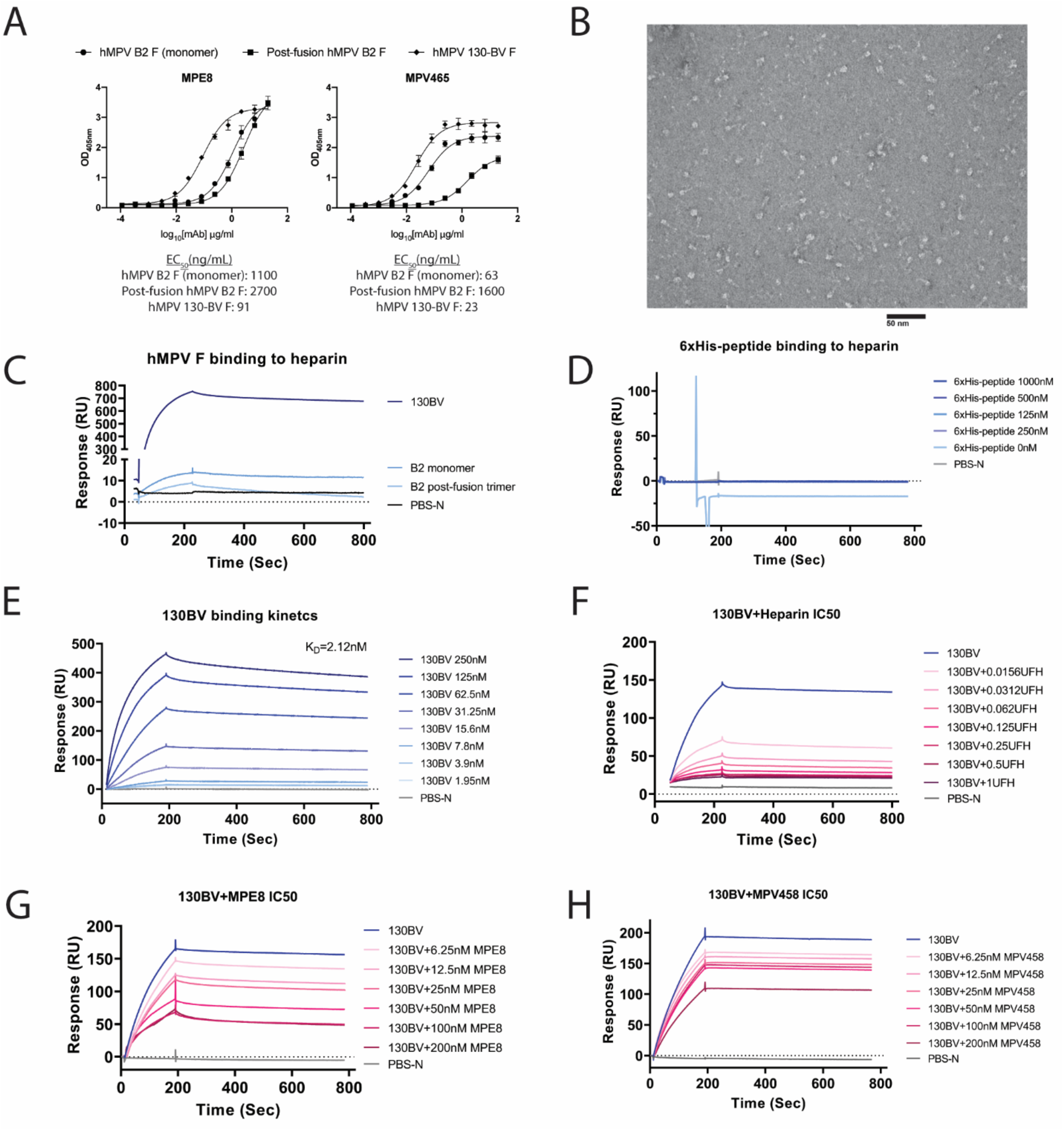
Recombinant hMPV F protein binds to heparin. (A) ELISA binding curves of MPE8 and MPV465 against hMPV B2 F monomer, hMPV B2 F post-fusion trimer, and hMPV 130-BV F. (B) Negative-stain electron micrograph of hMPV 130-BV F protein. (C) Surface Plasmon Resonance (SPR) curves hMPV B2 F monomer, hMPV B2 F post-fusion trimer, and hMPV 130-BV F binding to immobilized heparin. (D) Binding kinetics of 6×His-peptide to immobilized heparin. (E) Binding kinetics of hMPV 130-BV F at different concentrations (1.95nM – 250nM) to immobilized heparin. Concentration-dependent inhibition of hMPV 130-BV F binding to immobilized heparin with unfractionated heparin (F), MPE8 (G), and MPV458 (H).

To further determine the motif on HS that is required for hMPV F binding, we tested binding of hMPV F to a panel of oligomers of HS (36, 37). Similar to heparin, hMPV 130-BV showed overall stronger binding to the HS oligomers than monomeric or post-fusion hMPV B2 F. The microarray data showed hMPV F has a strong preference for specific HS oligosaccharides (compound 66, 78, 79, 91, and 92) **(Figure 4, Supplementary Table 1)**. All 5 compounds share the GlcNS6S-IdoA2S motif, indicating this unit potentially mediates the binding of HS to hMPV F. Since integrin α_5_β1 has also been shown to be a receptor for hMPV F (11, 12), we tested binding by ELISA and biolayer interferometry of multiple hMPV F constructs to recombinantly expressed integrin α_5_β1 using a stabilized construct as previously described (42), but we observed no binding of this protein to any recombinant hMPV F protein (data not shown). The lack of hMPV F binding to integrin α_5_β1 could be due to weak interactions, or the recombinant protein constructs may not be optimal for effective binding.

**Figure 4.**
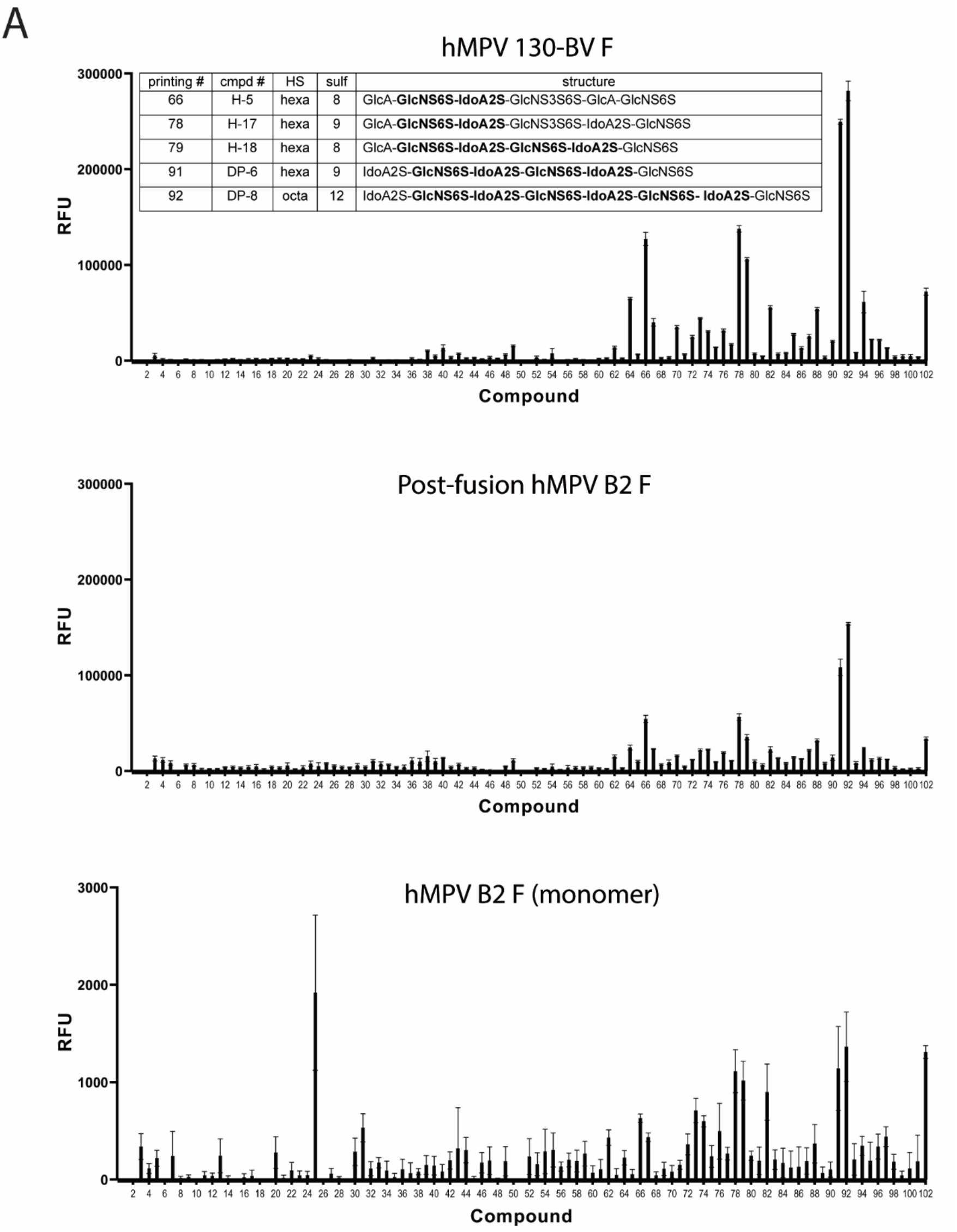
Heparan sulfate binding to hMPV F proteins by microarray. Binding of synthetic heparan sulfate oligosaccharides to hMPV 130-BV, post-fusion and monomeric hMPV B2 F proteins. The structures of the top 5 compounds for binding to hMPV 130-BV F protein are listed in the table.

### Vaccination with post-fusion hMPV B2 F induces a balanced immune response targeting two major epitopes

While the pre-fusion conformation of RSV F elicits more robust neutralizing antibody titers due to the presence of pre-fusion-specific antigenic sites (18), the majority of antigenic sites on hMPV F are present in both pre-fusion and post-fusion conformations (15), suggesting the more stable post-fusion F may be a viable vaccine candidate. However, there have been no studies to determine the protective efficacy of homogeneous post-fusion hMPV F. hMPV challenge studies following hMPV F protein vaccination have recently been reported, however, all constructs in that study were mixtures of pre-fusion and post-fusion conformations (17). To determine the immune response to vaccination with post-fusion hMPV F, mice were immunized with post-fusion hMPV F protein or PBS using a prime-boost-boost regimen as shown in **Figure 5A** using Titermax Gold adjuvant. The serum from post-fusion hMPV B2 F protein immunized mice showed increasing hMPV F-specific IgG levels after each immunization compared to pre-vaccination titers and the PBS+adjuvant immunized group **(Figure 5B)**. Immunized mice serum showed potent neutralization against both subgroups A2 (strain hMPV CAN/97-83) and B2 (strain hMPV TN/93-32) **(Figure 5C)**. In addition, A balanced Th1/Th2 immune response was observed as both robust IgG1 and IgG2a/2b were generated after the second boost **(Figure 5D)**. In all analyses, no major differences were observed between male and female groups of mice.

**Figure 5.**
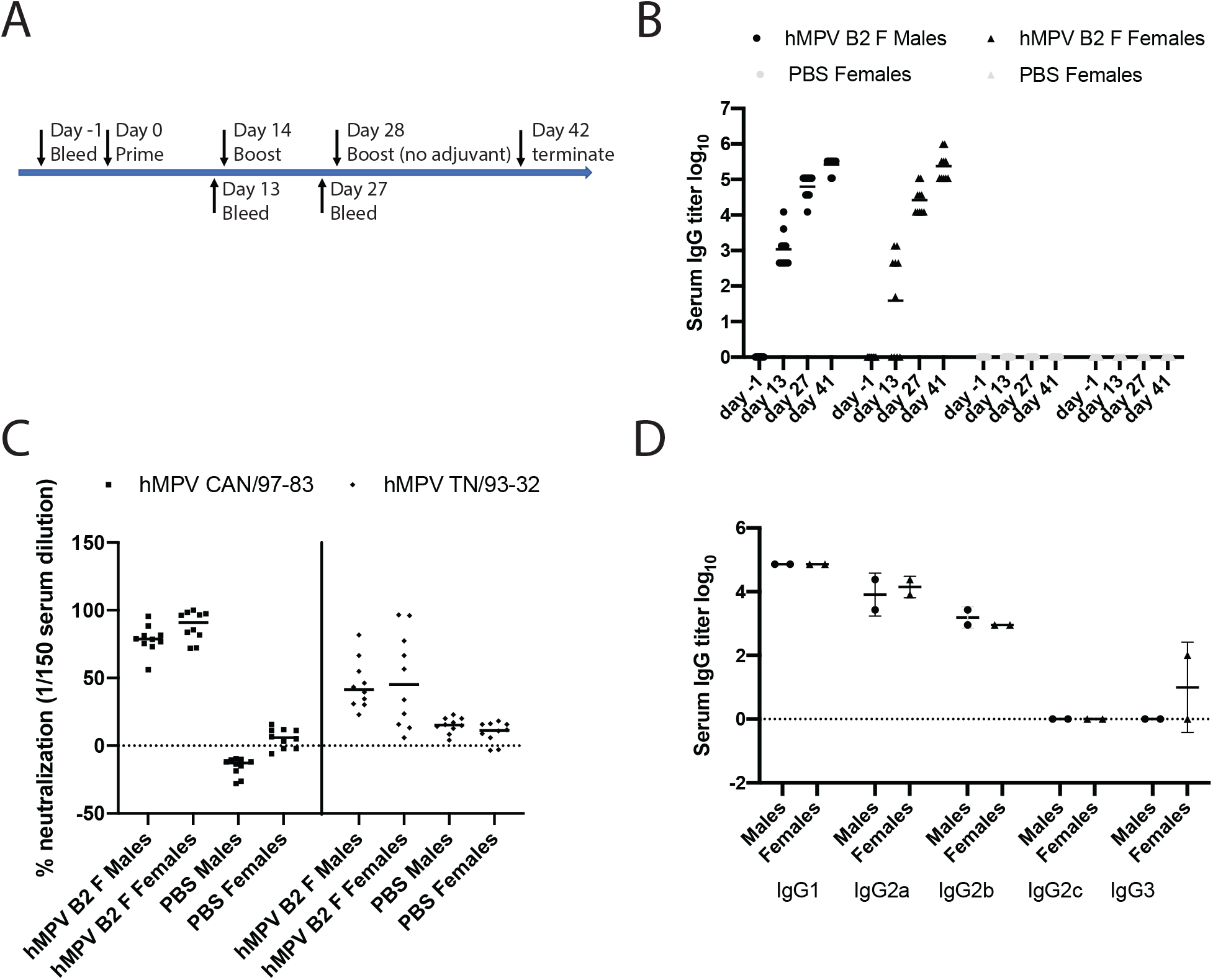
Mouse vaccination study with post-fusion hMPV B2 F. (A) Regimen of the vaccination study. (B) Serum hMPV F-specific IgG titers pre (day -1) and post (day 14, 28, 41) vaccinations. (C) Percent neutralization of 1/150 diluted endpoint serum against hMPV A2 (CAN/97-83) and B2 (TN/93-32). (D) Endpoint IgG subclasses titers against hMPV B2 F. Each data point represents a pool of serum samples from 5 mice in the same group.

To identify which antigenic sites were primarily targeted by post-fusion hMPV F immunization, monoclonal antibodies (mAbs) MPE8 (site III), 101F (site IV), DS7, and MPV458 (amino acids 66-87) were used in competition enzyme-linked immunosorbent assays with mouse serum against both monomeric (pre-fusion) and trimeric (post-fusion) hMPV F. Anti-human secondary antibody was used to detect mAb binding to recombinant proteins, while anti-mouse secondary antibody was used to detect the binding of serum mouse antibodies. To confirm the conformation of the proteins, the pre-fusion-specific mAb MPE8 was used, and this mAb showed binding to the monomeric (pre-fusion) F protein, but had limited binding to the trimeric (post-fusion) hMPV F protein. In both male (Groups 1-2) and female (Groups 3-4) mice, mAb competition with mouse serum was observed for mAbs MPE8 and 101F, and no competition was observed for mAbs DS7 and 458 **(Figure 6A)**. These data suggest that antigenic sites III and IV are predominantly targeted by mouse B cells in response to post-fusion hMPV F vaccination (**Figure 6B, 6C**).

**Figure 6.**
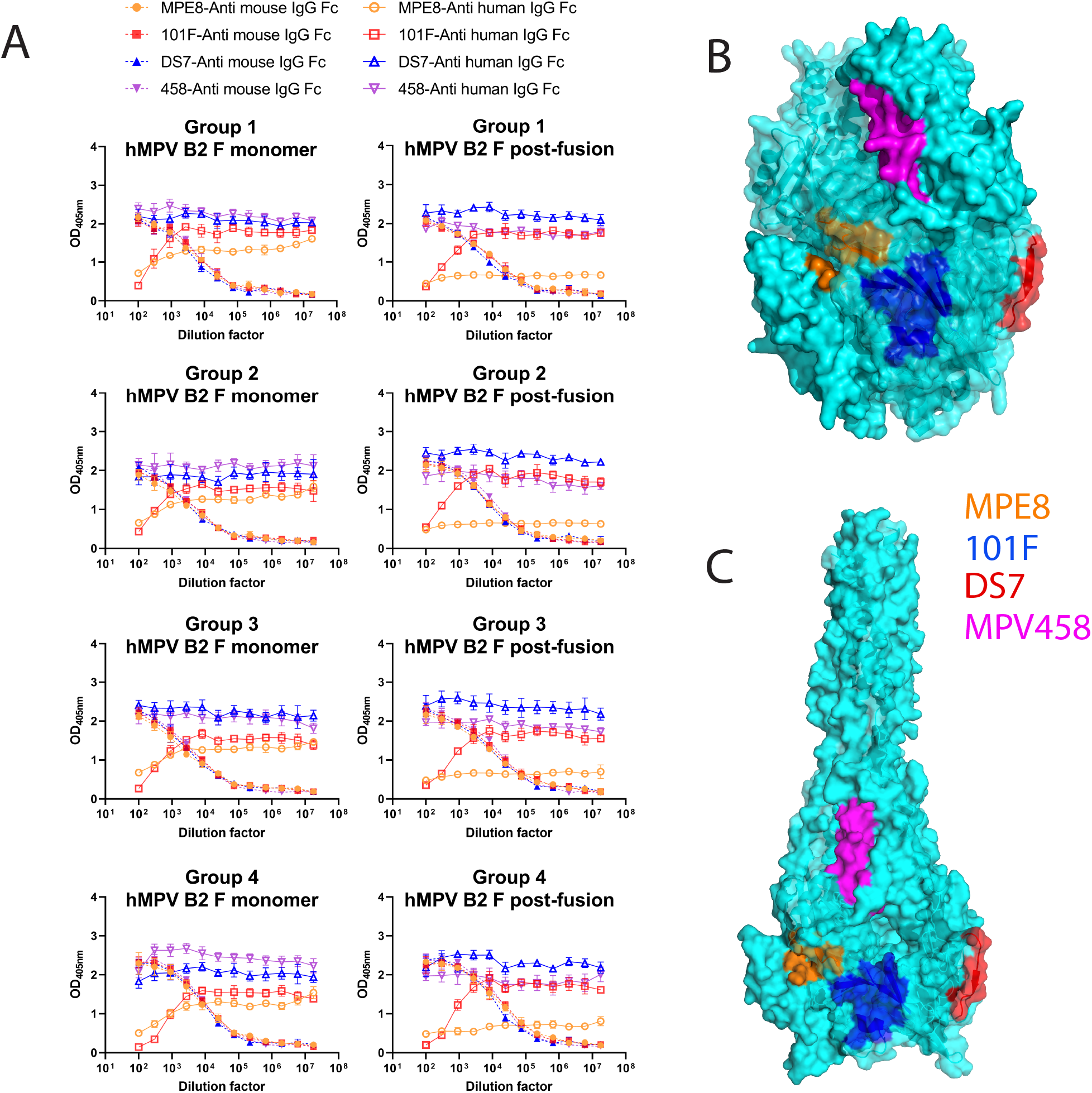
Competition ELISA of post-fusion hMPV B2 F vaccinated serum against human hMPV F mAbs. (A) Serial dilutions of vaccinated endpoint serum pooled from 5 mice in the same group competing with 4 human mAbs (MPE8-orange, 101F-red, DS7-blue, and MPV458-green). Mouse IgG was shown in dashed lines with solid symbols, while human IgG was shown in solid lines and empty symbols. Male mice are Groups 1-2, and female mice are Groups 3-4. The mAb binding sites of MPE8, 101F, DS7, and MPV458 are displayed on the surface of pre-fusion (B) and post-fusion (C) hMPV B2 F.

### Vaccination with post-fusion hMPV B2 limited viral replication

As vaccination with post-fusion hMPV B2 F elicited a robust and neutralizing IgG response, we next determined if such vaccination can protect against viral replication. In this study, mice were primed and boosted with post-fusion hMPV B2 F/PBS + TiterMax Gold adjuvant, then intranasally challenged with 5×10^5^ pfu hMPV B2 TN/93-32 two weeks after the boost **(Figure 7A)**. Vaccination with post-fusion hMPV F limited viral replication below the detection limit for all mice, as no virus was detected by the plaque assay in the lung homogenates **(Figure 7B)**. Vaccine enhanced disease is a potential concern with both RSV and hMPV, as formalin inactivation has previously been shown to exacerbate pulmonary pathological changes after challenge. We sectioned the lungs of two mice from each group, and scored peribronchiolotis, perivasculitis, interstitial pneumonitis, and alveolitis as previously described (23). Mild to moderate pathological changes were observed in hMPV F and PBS vaccinated groups compared to uninfected mice **(Figure 7D)**.

**Figure 7.**
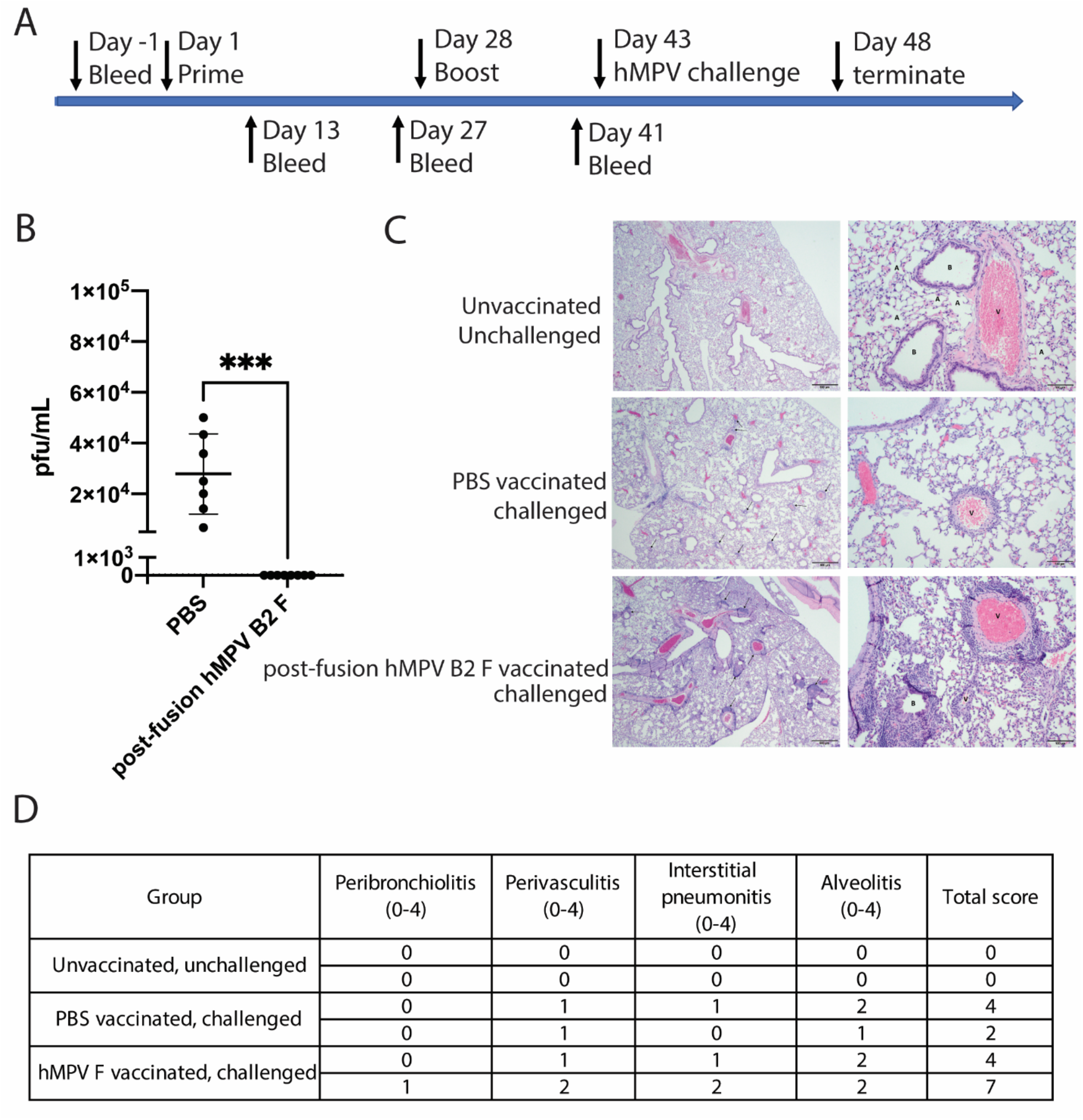
Mouse vaccination and challenge study with post-fusion hMPV B2 F. (A) Regimen of the vaccination and challenge study. (B) Endpoint lung viral loads were quantified by plaque forming units after immunostaining. Significant differences between unvaccinated and vaccinated groups are calculated by One-Way ANOVA, ***P=0.002 via unpaired t-test. (C) Represented figures show the lung histopathology of untreated mice and vaccinated mice at 2X magnification (left panel, scale bar indicates 500 μm) and 10X magnification (right panel, scale bar indicates 100 μm). (D) Pulmonary pathology changes were quantified by scores of peribronchiolitis, perivasculitis, interstitial pneumonitis, and alveolitis.

## Discussion

In this study, we determined the X-ray crystal structure of post-fusion hMPV F from genotype B, and determined the receptor binding properties, immunogenicity, and protective efficacy of this homogeneous recombinant protein. The X-ray structure of post-fusion hMPV B2 F closely aligns with the previously reported structure of post-fusion hMPV A1 F (16). While the hMPV A1 F protein was expressed in CV-1 cells using a vaccinia virus expression system (16), the hMPV B2 F could be expressed in milligram quantities from HEK293F cells. It is worth noting that the Jardetzky group has previously demonstrated the hMPV B2 F protein can be observed in the post-fusion conformation from HEK293F cells, although this protein contained a native cleavage and no high resolution structure of the protein in the post-fusion conformation was obtained (43). Overall, this expression protocol is reproducible and results in homogenous post-fusion hMPV F particles.

Both integrin α_5_β1 and heparan sulfate have been previously reported to be potential receptors for the hMPV F protein (8, 9, 12, 13). We demonstrated binding of heparin to both post-fusion hMPV B2 F and the pre-fusion-like protein hMPV 130-BV F. The pre-fusion conformation had much greater affinity for heparin than the post-fusion conformation. Furthermore, we determined the hMPV 130-BV binding to heparin is dependent on the concentration of F protein. We also localized heparin binding to antigenic sites III and 66-87 using hMPV F neutralizing mAbs MPE8 and MPV458 (30, 40). Finally, we determined the optimal heparan sulfate oligomers that bind to the hMPV F protein.

It has previously been reported that the majority of neutralizing human antibodies in serum target both pre-fusion and post-fusion conformations of the hMPV F protein, which suggests the post-fusion hMPV F protein may contain the majority of neutralizing antigenic sites (15). Based on these findings, we hypothesized that the immunodominant epitopes on hMPV F are conserved in both pre-fusion and post-fusion conformations. Therefore, immunization with post-fusion F would inducing antibodies that bind to pre-fusion F, which are capable of neutralizing the virus before fusion to host cells. To test this hypothesis, we vaccinated mice with post-fusion hMPV B2 F protein and determined that such vaccination elicits neutralizing antibodies that primarily target antigenic sites III and IV. Antigenic site IV is conformationally conserved in both pre-fusion and post-fusion conformations of hMPV F (44, 45). Antigenic site III elicits both pre-fusion-specific antibodies, such as MPE8, as well as antibodies that bind both pre-fusion and post-fusion conformations (29, 41).

Previous studies have implicated multiple factors that can be attributed to the FI-RSV vaccine-enhanced disease. The post-fusion conformation of the RSV F protein is dominate on FI-RSV particles (19), and such a vaccine cannot induce neutralizing antibodies to pre-fusion specific antigenic sites on the RSV F protein (46, 47). FI-RSV induces a Th2-skewed immune response that leads to eosinophil infiltration in the lungs, which may also contribute to the enhancement of the disease (48, 49). Similarly, FI-hMPV also boosts Th2 cytokines, however, neutralizing antibodies can still be induced to limit viral replication after challenge (22, 23). In this study, we found that post-fusion hMPV B2 F elicits a balanced Th1/Th2 immune response when used with Titermax adjuvant, which possibly alleviates the severity of pathological changes in the lungs. We then tested the protective efficacy of the hMPV B2 F protein and showed that vaccination completely protected mice from lung viral replication, consistent with result of the previous vaccination-challenge study using hMPV F proteins subtype A, which contained mixtures of pre-fusion and post-fusion conformations (17).

In summary, this work confirms the binding of hMPV F with its potential host receptor, heparan sulfate. The immunization and challenge studies bolster the potential of post-fusion hMPV F to become an effective vaccine candidate. Overall, these findings will shed light on the development of novel drugs and vaccines against hMPV.

## Acknowledgments

X-ray data were collected at the Southeast Regional Collaborative Access Team (SER-CAT) 22-ID beamLine at the Advanced Photon Source, Argonne National Laboratory. SER-CAT is supported by its member institutions (see www.ser-cat.org/members.htmL), and equipment grants (S10_RR25528 and S10_RR028976) from the National Institutes of Health. Use of the Advanced Photon Source was supported by the U.S. Department of Energy, Office of Science, Office of Basic Energy Sciences, under Contract No. W-31-109-Eng-38. We thank Georgia Electron Microscopy at the University of Georgia for assistance with negative-stain electron microscopy. The content is solely the responsibility of the authors and does not necessarily represent the official views of the National Institutes of Health. The structure factors and structure coordinates were deposited to the Protein Data Bank under accession code 7M0I. We thank Dr. John Williams (University of Pittsburgh) for helpful discussions on the mouse challenge viruses.

## Funding Statement

These studies were supported by National Institutes of Health grants 1R01AI143865 (JJM), 1K01OD026569 (JJM), and 5R01HL151617 (G-J.B.). The funders had no role in study design, data collection and analysis, decision to publish, or preparation of the manuscript.

**Supplementary Table.1.** Heparan sulfate oligosaccharides tested in this study.

| printing # | array<br>cmpd #<br>in<br>JACS<br>2017 | synthesis<br>cmpd #<br>in JACS<br>2017 | HS | sulf | backbone | structure |
| --- | --- | --- | --- | --- | --- | --- |
| 3 | 5-B | 53 | tetra | 5 | GIA-IdoA | GlcA-GlcNS6S-IdoA2S-GlcNS6S |
| 4 | 4-G | 34 | tetra | 4 | IdoA-GIA | IdoA-GlcNS6S-GlcA-GlcNS6S |
| 5 | 3-F | -<sup>1</sup> | tetra | 3 | GIA-IdoA | GlcA-GlcNAc6S-IdoA2S-GlcNAc6S |
| 6 |  |  | hexa | 6 |  | GlcA-GlcNS6S-IdoA-GlcNS6S-GlcA-GlcNS6S |
| 7 | 2-J | -<sup>1</sup> | tetra | 2 | IdoA-IdoA | IdoA-GlcNAc6S-IdoA-GlcNAc6S |
| 8 | 4-E | -<sup>1</sup> | tetra | 4 | GIA-IdoA | GlcA-GlcNS6S-IdoA-GlcNS6S |
| 9 | 2-H | 33 | tetra | 2 | IdoA-GIA | IdoA-GlcNAc6S-GlcA-GlcNAc6S |
| 10 | 2-F | -<sup>1</sup> | tetra | 2 | GIA-IdoA | GlcA-GlcNAc6S-IdoA-GlcNAc6S |
| 11 | 3-K | S34 | tetra | 3 | IdoA-IdoA | IdoA-GlcNAc6S-IdoA2S-GlcNAc6S |
| 12 | 4-B | -<sup>1</sup> | tetra | 4 | GIA-GIA | GlcA-GlcNS6S-GlcA-GlcNS6S |
| 13 | 4-I | 51 / S41 | tetra | 4 | IdoA-IdoA | IdoA-GlcNS6S-IdoA-GlcNS6S |
| 14 | 2-C | -<sup>1</sup> | tetra | 2 | GIA-GIA | GlcA-GlcNAc6S-GlcA-GlcNAc6S |
| 15 | 1-B | S14 | tetra | 1 | GIA-IdoA | GlcA-GlcNAc-IdoA2S-GlcNAc |
| 16 | 3-B | 47 / S15 | tetra | 3 | GIA-IdoA | GlcA-GlcNS-IdoA2S-GlcNS |
| 17 | 2-B | S22 | tetra | 2 | GIA-GIA | GlcA-GlcNAc-GlcA2S-GlcNAc6S |
| 18 | 4-A | S23 | tetra | 4 | GIA-GIA | GlcA-GlcNS-GlcA2S-GlcNS6S |
| 19 | 1-A | S20 | tetra | 1 | GIA-GIA | GlcA-GlcNAc-GlcA2S-GlcNAc |
| 20 | 3-A | S30 | tetra | 3 | GIA-GIA | GlcA-GlcNS-GlcA2S-GlcNS |
| 21 |  |  | tetra | 0 | GIA-GIA | GlcA-GlcNAc-GlcA-GlcNAc |
| 22 | 2-D | S10 | tetra | 2 | GIA-IdoA | GlcA-GlcNAc-IdoA-GlcNS6S |
| 23 | 4-D | 49 / S6 | tetra | 4 | GIA-IdoA | GlcA-GlcNS-IdoA2S-GlcNS6S |
| 24 | 2-E | S4 | tetra | 2 | GIA-IdoA | GlcA-GlcNAc-IdoA2S-GlcNAc6S |
| 25 | 3-D | S5 | tetra | 3 | GIA-IdoA | GlcA-GlcNAc-IdoA2S-GlcNS6S |
| 26 | 1-C | S9 | tetra | 1 | GIA-IdoA | GlcA-GlcNAc-IdoA-GlcNAc6S |
| 27 | 3-C | S11 | tetra | 3 | GIA-IdoA | GlcA-GlcNS-IdoA-GlcNS6S |
| 28 |  |  | tetra | 0 | GIA-IdoA | GlcA-GlcNAc-IdoA-GlcNAc |
| 29 |  |  | di | 1 |  | IdoA-GlcNAc6S |
| 30 | 1-E | 31 | tetra | 1 | IdoA-GIA | IdoA-GlcNAc6S-GlcA-GlcNAc |
| 31 | 5-C | 26 | tetra | 5 | IdoA-GIA | IdoA2S-GlcNS6S-GlcA-GlcNS6S |
| 32 | 3-H | 32 | tetra | 3 | IdoA-GIA | IdoA-GlcNS6S-GlcA-GlcNS |
| 33 | 3-I | 25 | tetra | 3 | IdoA-GIA | IdoA2S-GlcNAc6S-GlcA-GlcNAc6S |
| 34 | 2-G | 41 | tetra | 2 | IdoA-GIA | IdoA2S-GlcNAc6S-GlcA-GlcNAc |
| 35 | 1-D | 39 | tetra | 1 | IdoA-GIA | IdoA2S-GlcNAc-GlcA-GlcNAc |
| 36 | 4-F | 42 | tetra | 4 | IdoA-GIA | IdoA2S-GlcNS6S-GlcA-GlcNS |
| 37 | 3-G | 40 | tetra | 3 | IdoA-GIA | IdoA2S-GlcNS-GlcA-GlcNS |
| 38 | 5-D | S35 | tetra | 5 | IdoA-IdoA | IdoA-GlcNS6S-IdoA2S-GlcNS6S |
| 39 | 4-K | -<sup>1</sup> | tetra | 4 | IdoA-IdoA | IdoA2S-GlcNAc6S-IdoA2S-GlcNAc6S |
| 40 | 6-A | -<sup>1</sup> | tetra | 6 | IdoA-IdoA | IdoA2S-GlcNS6S-IdoA2S-GlcNS6S |
| 41 |  |  | hexa | 2 |  | GlcA-GlcNAc-IdoA2S-GlcNAc6S-GlcA-GlcNAc |
| 42 |  |  | hexa | 5 |  | GlcA-GlcNS-IdoA2S-GlcNS6S-GlcA-GlcNS |
| 43 |  |  | di | 2 |  | IdoA2S-GlcNAc6S |
| 44 |  |  | di | 3 |  | IdoA2S-GlcNS6S |
| 45 |  |  | di | 1 |  | GlcA-GlcNAc6S |
| 46 | CS-B | -<sup>2</sup> | tetra-CS | 2 |  | GlcA-GlcNS6S-GlcA-GlcNAc |
| 47 | 1-F | S40 | tetra | 1 | IdoA-IdoA | IdoA-GlcNAc-IdoA-GlcNAc6S |
| 48 | 1-G | S39 | tetra | 1 | IdoA-IdoA | IdoA-GlcNAc6S-IdoA-GlcNAc |
| 49 |  |  | tetra | 6 | GIA-IdoA | GlcA-GlcNS3S6S-IdoA2S-GlcNS6S |
| 50 | 5-E | 52 | tetra | 5 | IdoA-IdoA | IdoA-GlcNS3S6S-IdoA-GlcNS6S |
| 51 | 4-C | 48 | tetra | 4 | GIA-IdoA | GlcA-GlcNS3S-IdoA2S-GlcNS |
| 52 | 5-A | 50 | tetra | 5 | GIA-IdoA | GlcA-GlcNS3S-IdoA2S-GlcNS6S |
| 53 | CS-A | -<sup>2</sup> | tetra-CS | 1 |  | GlcA-GalNAc-GlcA-GalNAc4S |
| 54 |  |  | octa | 7 |  | GlcA-GlcNS6S-IdoA-GlcNS-IdoA2S-GlcNS6S-IdoA-GlcNAc6S |
| 55 | 2-A | S51 | tetra | 2 | GIA-GIA | GlcA-GlcNS6S-GlcA-GlcNAc |
| 56 | 2-I | S43 | tetra | 2 | IdoA-IdoA | IdoA-GlcNS-IdoA-GlcNS |
| 57 | 4-H | S45 | tetra | 4 | IdoA-IdoA | IdoA-GlcNS-IdoA2S-GlcNS6S |
| 58 | 3-E | S53 | tetra | 3 | GIA-IdoA | GlcA-GlcNS6S-IdoA-GlcNS |
| 59 | 4-J | S47 | tetra | 4 | IdoA-IdoA | IdoA2S-GlcNS6S-IdoA-GlcNAc6S |
| 60 | 3-J | S49 | tetra | 3 | IdoA-IdoA | IdoA-GlcNS-IdoA-GlcNS6S |

| printing # | compd # | HS | sulf | structure |
| --- | --- | --- | --- | --- |
| 62 | H-1 | hexa | 7 | GlcA-GlcNS6S-GlcA-GlcNS3S6S-GlcA-GlcNS6S |
| 63 | H-2 | hexa | 6 | GlcA-GlcNS6S-GlcA-GlcNS6S-GlcA-GlcNS6S |
| 64 | H-3 | hexa | 7 | GlcA-GlcNS6S-IdoA-GlcNS3S6S-GlcA-GlcNS6S |
| 65 | H-4 | hexa | 6 | GlcA-GlcNS6S-IdoA-GlcNS6S-GlcA-GlcNS6S |
| 66 | H-5 | hexa | 8 | GlcA-GlcNS6S-IdoA2S-GlcNS3S6S-GlcA-GlcNS6S |
| 67 | H-6 | hexa | 7 | GlcA-GlcNS6S-IdoA2S-GlcNS6S-GlcA-GlcNS6S |
| 68 | H-7 | hexa | 7 | GlcA-GlcNS6S-GlcA-GlcNS3S6S-IdoA-GlcNS6S |
| 69 | H-8 | hexa | 6 | GlcA-GlcNS6S-GlcA-GlcNS6S-IdoA-GlcNS6S |
| 70 | H-9 | hexa | 7 | GlcA-GlcNS6S-IdoA-GlcNS3S6S-IdoA-GlcNS6S |
| 71 | H-10 | hexa | 6 | GlcA-GlcNS6S-IdoA-GlcNS6S-IdoA-GlcNS6S |
| 72 | H-11 | hexa | 8 | GlcA-GlcNS6S-IdoA2S-GlcNS3S6S-IdoA-GlcNS6S |
| 73 | H-12 | hexa | 7 | GlcA-GlcNS6S-IdoA2S-GlcNS6S-IdoA-GlcNS6S |
| 74 | H-13 | hexa | 8 | GlcA-GlcNS6S-GlcA-GlcNS3S6S-IdoA2S-GlcNS6S |
| 75 | H-14 | hexa | 7 | GlcA-GlcNS6S-GlcA-GlcNS6S-IdoA2S-GlcNS6S |
| 76 | H-15 | hexa | 8 | GlcA-GlcNS6S-IdoA-GlcNS3S6S-IdoA2S-GlcNS6S |
| 77 | H-16 | hexa | 7 | GlcA-GlcNS6S-IdoA-GlcNS6S-IdoA2S-GlcNS6S |
| 78 | H-17 | hexa | 9 | GlcA-GlcNS6S-IdoA2S-GlcNS3S6S-IdoA2S-GlcNS6S |
| 79 | H-18 | hexa | 8 | GlcA-GlcNS6S-IdoA2S-GlcNS6S-IdoA2S-GlcNS6S |
| 80 | H-19 | hexa | 6 | GlcA-GlcNS6S-GlcA-GlcNS3S-GlcA-GlcNS6S |
| 81 | H-20 | hexa | 6 | GlcA-GlcNS6S-IdoA-GlcNS3S-GlcA-GlcNS6S |
| 82 | H-21 | hexa | 7 | GlcA-GlcNS6S-IdoA2S-GlcNS3S-GlcA-GlcNS6S |
| 83 | H-22 | hexa | 6 | GlcA-GlcNS6S-GlcA-GlcNS3S-IdoA-GlcNS6S |
| 84 | H-23 | hexa | 6 | GlcA-GlcNS6S-IdoA-GlcNS3S-IdoA-GlcNS6S |
| 85 | H-24 | hexa | 7 | GlcA-GlcNS6S-IdoA2S-GlcNS3S-IdoA-GlcNS6S |
| 86 | H-25 | hexa | 6 | GlcA-GlcNS6S-GlcA-GlcNS3S-IdoA2S-GlcNS6S |
| 87 | H-26 | hexa | 7 | GlcA-GlcNS6S-IdoA-GlcNS3S-IdoA2S-GlcNS6S |
| 88 | H-27 | hexa | 8 | GlcA-GlcNS6S-IdoA2S-GlcNS3S-IdoA2S-GlcNS6S |
| 89 | DP-2 | di | 3 | IdoA2S-GlcNS6S |
| 90 | DP-4 | tetra | 6 | IdoA2S-GlcNS6S-IdoA2S-GlcNS6S |
| 91 | DP-6 | hexa | 9 | IdoA2S-GlcNS6S-IdoA2S-GlcNS6S-IdoA2S-GlcNS6S |
| 92 | DP-8 | octa | 12 | IdoA2S-GlcNS6S-IdoA2S-GlcNS6S-IdoA2S-GlcNS6S-IdoA2S-GlcNS6S |
| 93 | SLF-134 | hexa | 5 | GlcA-GlcNAc-IdoA2S-GlcNS6S-IdoA2S-GlcNS |
| 94 | SLF-136 | hexa | 7 | GlcA-GlcNS6S-IdoA2S-GlcNS6S-IdoA2S-GlcNS |
| 95 | SLF-139 | hexa | 6 | GlcA-GlcNS-IdoA2S-GlcNS6S-IdoA2S-GlcNS |
| 96 | SLF-140 | hexa | 6 | GlcA-GlcNAc6S-IdoA2S-GlcNS6S-IdoA2S-GlcNS |
| 97 | SLF-138 | tetra | 5 | IdoA2S-GlcNS6S-IdoA2S-GlcNS |
| 98 | PC-160 | tetra | 5 | GlcA-GlcNS6S-GlcA2S-GlcNS6S |
| 99 | PC-161 | tetra | 3 | GlcA-GlcNAc6S-GlcA2S-GlcNAc6S |
| 100 | PC-141 | tetra | 1 | IdoA-GlcNS-IdoA-GlcNAc |
| 101 | PC-142 | tetra | 1 | IdoA-GlcNAc-IdoA-GlcNS |
| 102 | Heparin amine | polymer | -- | Heparin functionalized with primary amine (MW 27Kda, CreativePEGworks) |

Abbreviations:
GlcA = D-glucuronic acid
GlcNAc = *N*-acetyl-D-glucosamine
GlcN = glucosamine
IdoA = L-iduronic acid
GalNAc = *N*-acetylgalactosamine

## Notes

### Competing Interest Statement

The authors have declared no competing interest.

